# Using Arabian Sea Coastal Water as a Model to Study Plasmid-mediated AMR Dissemination driven by Anthropogenic Impacts

**DOI:** 10.64898/2026.10.21.760660

**Authors:** Devika Ghadigaonkar, Devashish Rath, Archana Rath

## Abstract

**Background:** Antimicrobial resistance (AMR) is a growing global public health threat. The water bodies act as both reservoirs and medium for dissemination of AMR. Coastal water driven by anthropogenic effects accumulate antibiotics, heavy metals, microplastics, and disinfectants which act as stressors, increasing the overall AMR. Therefore, it is necessary to understand the AMR entities and its transmission in coastal water bodies.

**Methods and Results:** Mumbai, mirroring most cosmopolitan cities of the world, was used as a model to understand how rapid urbanization disseminate AMR in coastal waters of the Arabian Sea and can pose substantial risk to human health. Bacteria isolated from coastal waters were screened and found to be multi-drug resistant. Several isolates carried multiple plasmids of high molecular weight (>10 kb). Plasmid sequencing and analysis using various bioinformatic tools identified plasmid replicons like IncFIA, IncFII, rep7a and repUS43. Analysis showed a high burden of antibiotic resistance genes (ARGs) conferring resistance to antibiotic classes such as monobactams (bla_*TEM-116*_), carbapenems (bla_*NDM-5*_), vancomycin (*vanA*), disinfectants and antiseptics (*qacG*). Additionally, clinically relevant *mecA* and *mecR1* associated with methicillin-resistant *Staphylococcus* sp., *rpsL* associated with streptomycin resistant *Mycobacterium tuberculosis* and the colistin-resistance genes *mcr-10*.*1* and *mcr-10*.*4* were also detected. We also report for the first-time presence of a plasmid-borne *mupB* gene in coastal waters, that confers high resistance to mupirocin.

**Conclusions:** Despite the moderate sample size, the presence of diverse plasmid replicons and clinically important ARGs suggests urban coastal environment as a potent reservoir of AMR, with potential for global transmission through active waterway trade and transport.

## 1. Introduction

Antimicrobial resistance (AMR) is a rapidly growing public health threat, increasingly contributing to global disease burden and mortality. The emergence of multidrug-resistant (MDR), extensively drug-resistant (XDR), and pan drug-resistant (PDR) bacterial pathogens has rendered many first-line and even last-resort antibiotics ineffective, resulting in increased healthcare costs, treatment failures, and higher mortality rates [1].

Antibiotic-resistant bacteria (ARB) in the environment and the resistance factors they carry, like antibiotic resistance genes (ARGs) and mobile genetic elements (MGEs), can be transmitted to humans through food, drinking water, air, or through direct contact with the environment, such as through recreational activities in coastal waters [1]. Aquatic systems, by serving as reservoirs and transmission pathways for AMR, link environmental, clinical, and agricultural domains in the context of a One Health framework [2, 3, 4]. Studies show that coastal water containing antibiotics, heavy metals, microplastics, hydrocarbons, and disinfectants act as stressors, thus increasing the overall AMR of the environment [4, 5]. The negative effects of anthropogenic activities are particularly severe in the coastal zone of the Indian subcontinent due to the warn climate conditions [6]. As AMR from water bodies can get transmitted to humans via food chain [7, 8], it has profound implications for both ecosystem functioning and human health. Therefore, it is necessary to understand the AMR entities and its transmission in coastal water bodies.

Among the various mechanisms responsible for bacterial adaptation, horizontal gene transfer (HGT) plays a central role in the dissemination of AMR, with plasmids acting as efficient vectors. Plasmids often carry genes conferring resistance to multiple antibiotic classes, heavy metals, disinfectants, and virulence determinants, with high molecular weight plasmids (>10 kb) frequently having many of these genes on the same plasmid [9].

Mumbai was used as a model to study role of tropical urbanization and consequent anthropogenic effects on dissemination of AMR in coastal waters of the Arabian Sea.

Here, we performed a meta-analysis of plasmid DNA (pDNA) extracted from multi-ARB isolated from the coastal waters of Arabian Sea in Mumbai, India. The study identified diverse plasmid-associated ARGs, many of them linked to be originating from various anthropogenic activities like sewage and agriculture run-offs, and hospital waste discharge. By identifying various plasmid replicons, plasmid incompatibility groups and plasmid-associated ARGs, this study reiterates the role of plasmids in the environmental dissemination of antimicrobial resistance.

## 2. Materials and Methods

### 2.1 Sample Collection and Pre-processing

Two liters of samples were collected from two most visited coastal regions (Versova Beach and Juhu Beach) of the Arabian Sea in Mumbai, India, during March 2022 (pre-monsoon) and December 2022 (post monsoon; Table 1 and Fig 1). Sampling was done from a depth of approximately 0.5 m in sterile PET bottles, transported to the lab in ice boxes and processed immediately by filtering using sterile muslin cloth to remove any solid matter. These filtered samples were used for further analysis.

**Table 1.** Water sampling from two coastal regions of the Arabian Sea, Mumbai, India.

| Sample Code | Sampling Location | Sampling Season | GIS Coordinates |
| --- | --- | --- | --- |
| p1 | Versova Beach, Mumbai | Pre-monsoon | latitude 19.130383° N and longitude 72.812931° E |
| p2 | Juhu Beach, Mumbai | Pre-monsoon | latitude 19.10939° N and longitude 72.82407 ° E |
| p3 | Versova Beach, Mumbai | Post-monsoon | latitude 19.130383° N and longitude 72.812931° E |
| p4 | Juhu Beach, Mumbai | Post-monsoon | latitude 19.10939° N and longitude 72.82407 ° E<br>72.82° E |

**Fig. 1.**
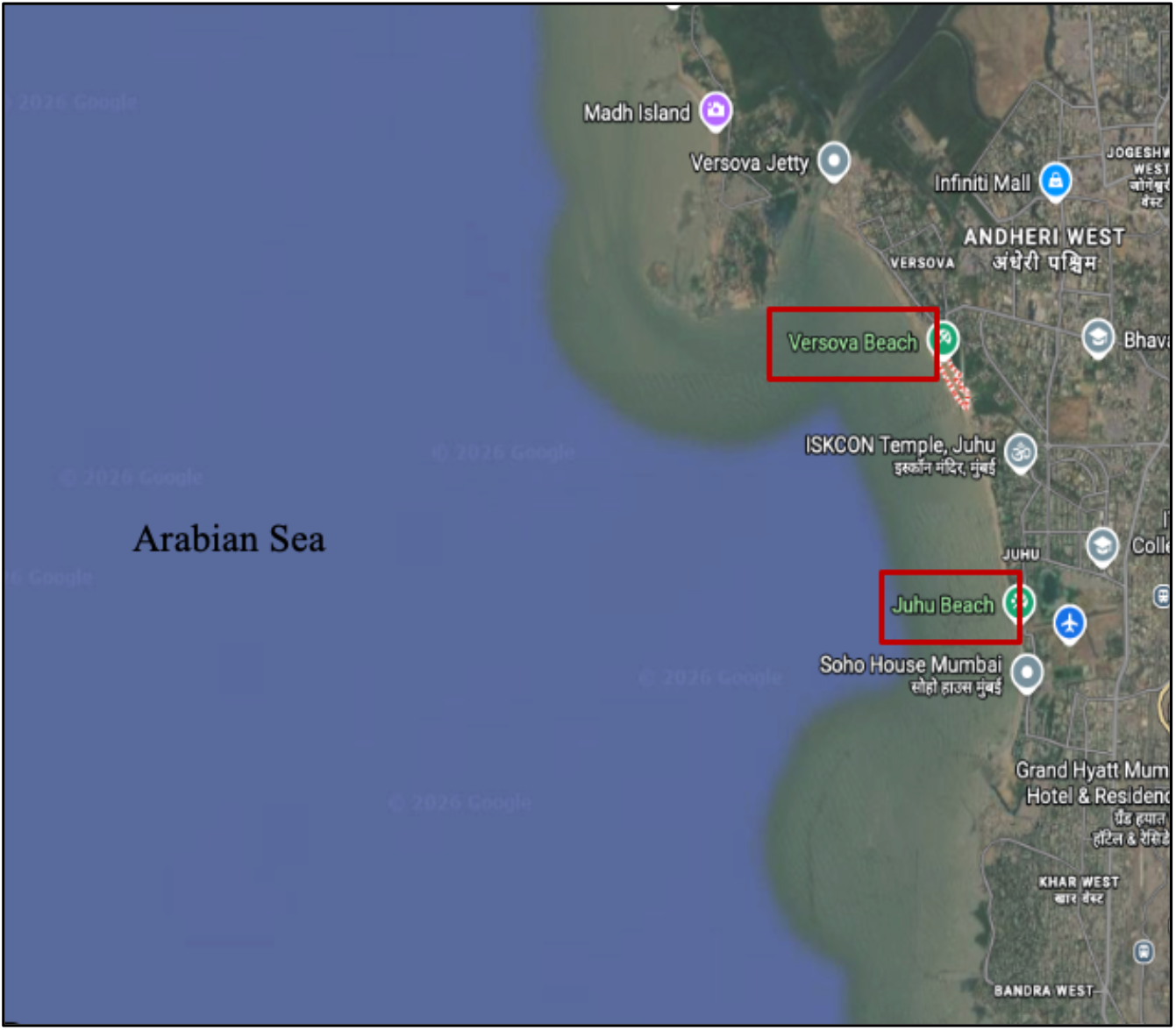
The map shows the two sampling points of the Arabian Sea in Mumbai, India

### 2.2 Isolation of antibiotic-resistant bacteria

Enumeration of total and antibiotic-resistant bacteria (ARB) was done using previously described method [10]. For primary isolation of ARB, filtered water samples were serially diluted using 0.85% saline and inoculated on Zobell Marine Agar media (HiMedia) containing individual antibiotics, namely cefotaxime (CTX, 3 µg/mL), ciprofloxacin (CIP, 0.5 µg/mL), erythromycin (E, 20 µg/mL), kanamycin (K, 15 µg/mL) and vancomycin (VA, 3 µg/mL), and incubated at 25 °C for 40-72 h. Post incubation, the total and ARB colony forming units per mL (CFU/mL) were calculated. Well-isolated ARB colonies were selected from each plate, purified for two passages on medium containing respective antibiotic and stored as glycerol stocks at −40 °C for subsequent study.

### 2.3 Identification of ARB using 16S rRNA gene sequencing

Twenty-five ARB from each sample were chosen for further study. Crude DNA extracted from bacterial lysate [10] was used to amplify the 457 bp internal fragment of the 16S rRNA gene using KAPA Taq Ready Mix, forward primer: 5′-GGAGGCAGCAGTAAGGAAT-3′ and reverse primer: 5′-CTACCGGGGTATCTAATCC-3′. These primers target the highly variable V3 and V4 regions of the 16S rRNA gene [11]. Reaction was set using 5 µL 2X Taq PCR Master Mix, 0.5 µL each of forward and reverse primers (stock concentration of 10 µM each), 1.5 µL of DNA. The volume was made up to 10 µL with sterile PCR grade water. The cycling condition used was: initial denaturation at 94 °C for 10 min followed by 35 cycles of denaturation at 94 °C for 30 s, annealing at 50 °C for 30 s and extension at 72 °C for 40 s. A final extension was carried out at 72 °C for 5 min. After confirmation by agarose gel electrophoresis on 1.5 % agarose, the amplicons were purified and subjected to Sanger sequencing. The sequence identity was checked by Basic Local Alignment Search Tool (BLAST) with non-redundant database.

### 2.4 Antibiotic Susceptibility Testing

Antibiotic Susceptibility Testing (AST) by disc diffusion method was carried out for the identified isolates using 12 clinically relevant antibiotics following CLSI (2015) [12] and EUCAST (2022) [13] standard protocols, as described earlier [10]. Antibiotics, namely amikacin (AK), azithromycin (AZM), aztreonam (AT), cefoxitin (CX), cefotaxime (CTX), ciprofloxacin (CIP), erythromycin (E), imipenem (IMP), levofloxacin (LE), tigecycline (TGC), trimethoprim (TR) and vancomycin (VA), were selected as per the WHO list of critically important antibiotics class (WHO 2018) [14]. The tests were performed in triplicates. *E. coli* ATCC 25922 and *S. aureus* ATCC 29213 were used as internal controls. Based on AST results, multiple antibiotic resistance (MAR) and multiple antibiotic resistance index (MARi), which is the ratio of number of antibiotics to which the isolate is resistant to the total number of antibiotics to which isolate is exposed, was calculated for each isolate.

### 2.5 Plasmid DNA extraction

Approximately ten isolates per sample per season, showing complete or intermediate resistance to at least two antibiotics were selected. pDNA from these ARB was extracted using Plasmid Extraction mini kit (Favorgen) following manufacturer’s instructions. pDNA concentration was quantified using the Synergy H1m Multimode Reader (BioTek) and plasmid profile analysed using agarose gel electrophoresis (AGE). pDNA from all the isolates which gave good quality plasmids were pooled together to obtain four pooled plasmid samples corresponding to each sample location and season (p1, p2, p3 & p4). pDNA samples were quantified using Qubit dsDNA High Sensitivity Assay (Invitrogen, Cat#Q32854) and purity checked on Multiskan Sky High Microplate Spectrophotometer. The samples were then processed for high throughput sequencing.

### 2.6 Library preparation and Sequencing

The libraries were constructed using the Twist Library Preparation Enzymatic Fragmentation Kit (TWIST Biosciences, Cat#: 104175). Library QC was carried out with 5300 Fragment Analyzer system (Agilent). Libraries were quantified using Qubit HS Assay (Invitrogen, Cat#Q32854) and sequenced in pair-end mode (2 × 150 bp) using the Illumina NovaSeq X plus platform. The raw data was submitted to the NCBI’s Sequence Read Archive (SRA, BioProject accession number PRJNA1422493).

### 2.7 Bioinformatics Analysis of sequence reads

#### Quality Control

Raw-read quality was checked using FastQC. Adapters were removed and low-quality reads (Phred score < 30) were filtered and reads with Q ≥ 30 were used for downstream analysis. The mean Phred score for all samples was found to be between 36 and 39.

#### Plasmid Assembly and Identification

For plasmid assembly, mpSPAdes (v. 4.3.0) was used (https://bio.tools/metaplasmidspades). The circular plasmid sequences obtained were submitted to NCBI’s GenBank (Accession numbers: PZ221391-PZ221420). Assembled sequences were further analysed using BLAST to check the identity of the plasmids. In addition, PlasmidFinder (v.2.0.1) (https://cge.food.dtu.dk/services/PlasmidFinder/) with minimum identity and coverage of 95% and 60% respectively was used to identify the plasmid replicons. Both databases Gram Positive and Enterobacteriales of PlasmidFinder 2.1 were used.

#### Resistome analysis

ARGs were identified using the antimicrobial resistance pipeline from CZ ID (formerly IDseq) (https://czid.org). This pipeline also generates contigs from the sequence reads to match them with the ARGs using the Resistance Gene Identifier tool from Comprehensive Antibiotic Resistance Database (RGI-CARD) (RGI 6.0.5, CARD 4.0.1) with the default selection criteria “perfect and strict hits only”. In addition to the CARD database, the contigs were also analyzed using ResFinder (v. 4.7.2).

A pictorial depiction of the methodology used is shown in Fig 2.

**Fig. 2.**
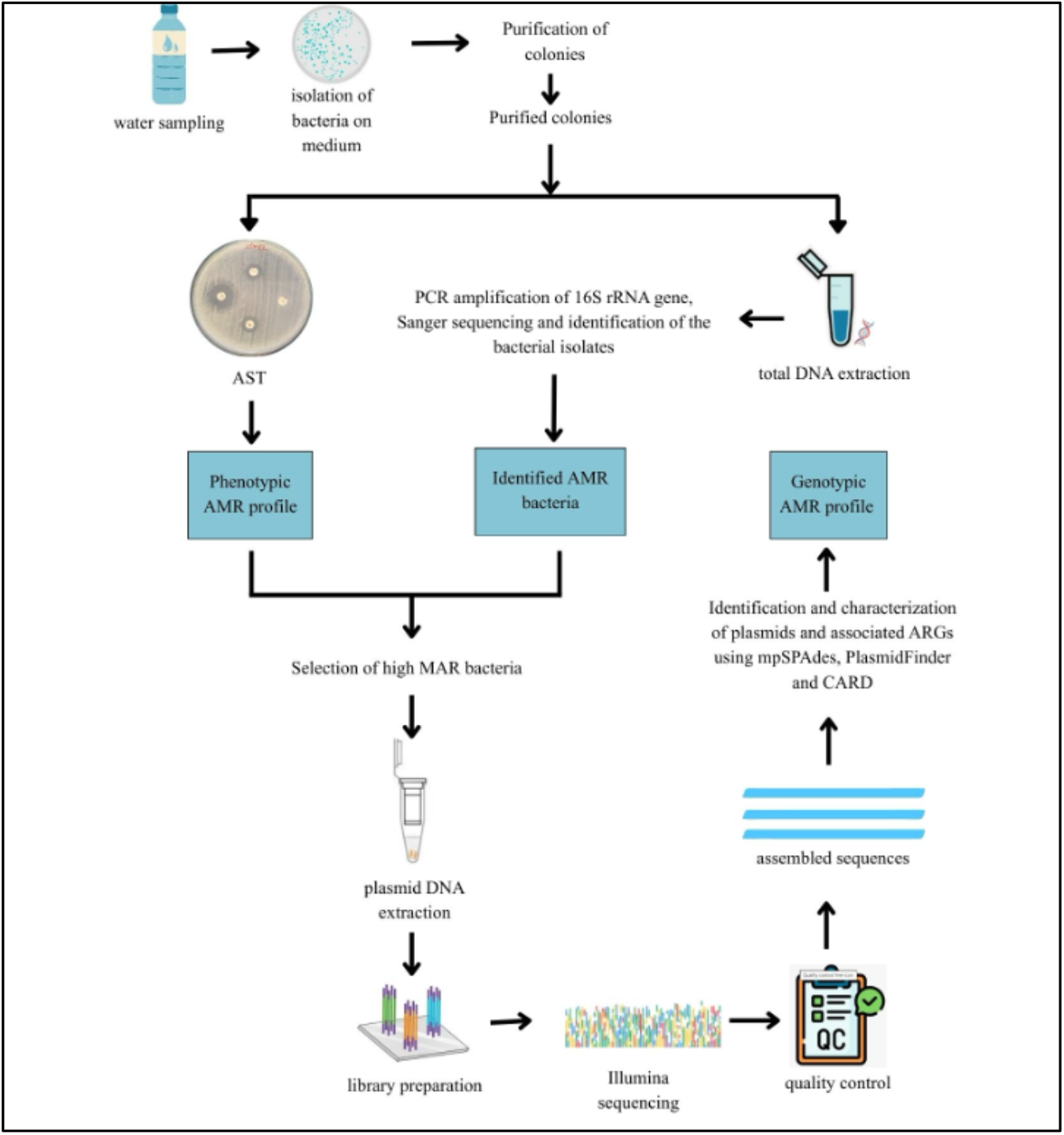
The study workflow shows the isolation of ARB, extraction of plasmid DNA from high MAR bacteria followed by high throughput Illumina sequencing and processing of reads for identification of plasmids, plasmid replicons and ARGs ARB: antibiotic resistant bacteria; MAR: multiple antibiotic resistant; ARGs: antibiotic resistance genes; AST: antibiotic susceptibility testing; AMR: antimicrobial resistance; mpSPAdes: metaplasmidSPAdes; CARD: The Comprehensive Antibiotic Resistance Database

## 3. Results

### 3.1 Prevalence of ARB in coastal water and their resistivity profile

In order to know the ARB load in water samples, total CFU (without antibiotic) vs antibiotic resistance (AR) CFU (with antibiotic) was calculated (Table 2). For pre-monsoon sample p1, the total CFU/mL was 1.6 × 10^5^ CFU/mL and the AR CFU/mL was between 1.5 × 10^3^ to 5.95 ×10^4^. For sample p2, the total CFU/mL was 7.1 × 10^5^ CFU/mL and the AR CFU/mL was between 1.0 × 10^4^ to 4 ×10^5^. For post-monsoon samples p3, despite serial dilutions, the total CFU/mL was found to be too numerous to count (TNTC) and the AR CFU/mL was between 8.0 × 10^6^ to 5.2 ×10^7^. For post-monsoon sample p4, the total CFU/mL was 8.65 × 10^8^ CFU/mL and the AR CFU/mL was between 4.2 × 10^4^ to 1.32 ×10^6^.

**Table 2.**
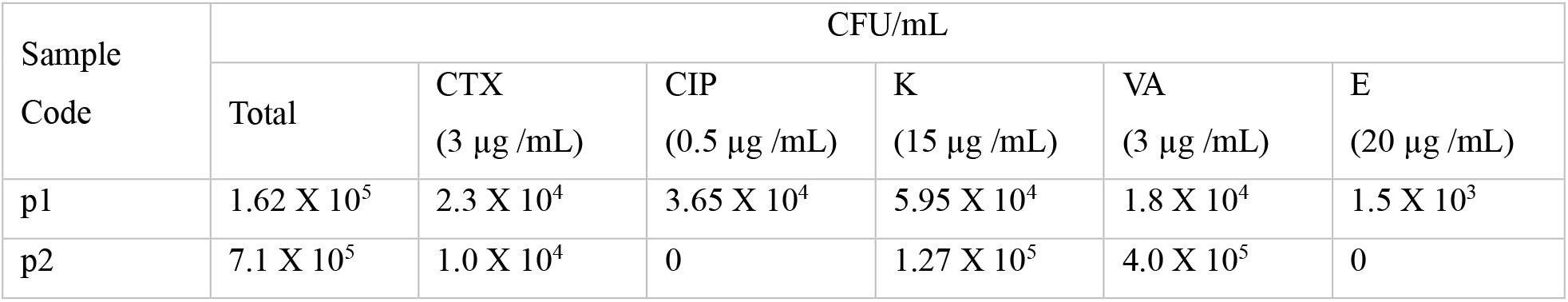
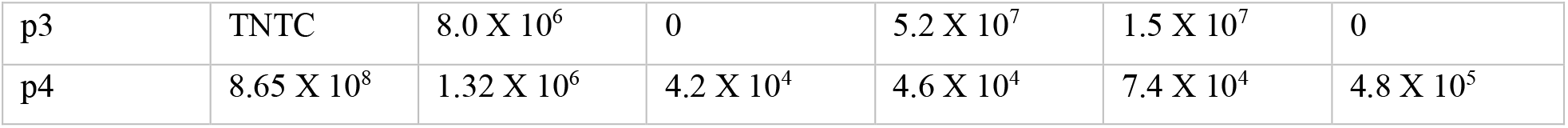
Total and ARB load in the pre-monsoon and post monsoon coastal water samples ARB were identified by 16S rRNA sequencing and subsequent BLASTn analysis. ARB predominantly belonged to *Enterobacteriaceae, Enterococcaceae, Staphylococcaceae, Bacillaceae*, and *Moraxellaceae*.

| Sample Code | CFU/mL |  |  |  |  |  |
| --- | --- | --- | --- | --- | --- | --- |
|  | Total | CTX<br>(3 µg /mL) | CIP<br>(0.5 µg /mL) | K<br>(15 µg /mL) | VA<br>(3 µg /mL) | E<br>(20 µg /mL) |
| p1 | $1.62 \times 10^5$ | $2.3 \times 10^4$ | $3.65 \times 10^4$ | $5.95 \times 10^4$ | $1.8 \times 10^4$ | $1.5 \times 10^3$ |
| p2 | $7.1 \times 10^5$ | $1.0 \times 10^4$ | 0 | $1.27 \times 10^5$ | $4.0 \times 10^5$ | 0 |
| p3 | TNTC | 8.0 X 10<sup>6</sup> | 0 | 5.2 X 10<sup>7</sup> | 1.5 X 10<sup>7</sup> | 0 |
| p4 | 8.65 X 10<sup>8</sup> | 1.32 X 10<sup>6</sup> | 4.2 X 10<sup>4</sup> | 4.6 X 10<sup>4</sup> | 7.4 X 10<sup>4</sup> | 4.8 X 10<sup>5</sup> |

AST profile using 12 clinically relevant antibiotics showed that most isolates were resistant to at least one or multiple antibiotics. The identity of the final selected ARB used for the present study, along with their AST profile is shown in the supplementary information (SI) Table S1 and Table S2 respectively.

In order to study plasmid mediated resistance, plasmids were extracted from these MAR isolates and analysed.

### 3.2 Plasmid profile of MAR isolates

pDNA extracted from MAR isolates were checked for the plasmid profile. Most of these isolates showed presence of more than one plasmid. Many isolates also contained one/ multiple high molecular weight (> 10 kb) plasmids. Since, such mega plasmids are known to harbor ARGs, their presence suggests that plasmids may be a major source of resistance acquisition in these isolates [9, 15]. The representative image of the plasmid profile of few AR isolates is shown in SI Fig S1.

### 3.3 Annotation of plasmids and plasmid replicons

High quality sequence reads assembled with mpSPAdes (v. 4.3.0) generated 30 circular plasmid sequences which were submitted to NCBI’s GenBank with accession numbers - PZ221391 to PZ221420 (SI Table S3). To identity the plasmids, the assembled sequences were analysed using BLAST. Multiple full-length plasmids were found to be present in each sample (Table 3). Using PlasmidFinder, various plasmid replicons were detected with an identity of more than 99%. (at the time of analysis, PlasmidFinder database contained 133 unique plasmid replicon sequences from members of Enterobacteriales and Gram-positive bacteria). The different types of plasmid families and plasmid replicons identified in each sample is summarized in Table 3.

**Table 3.**
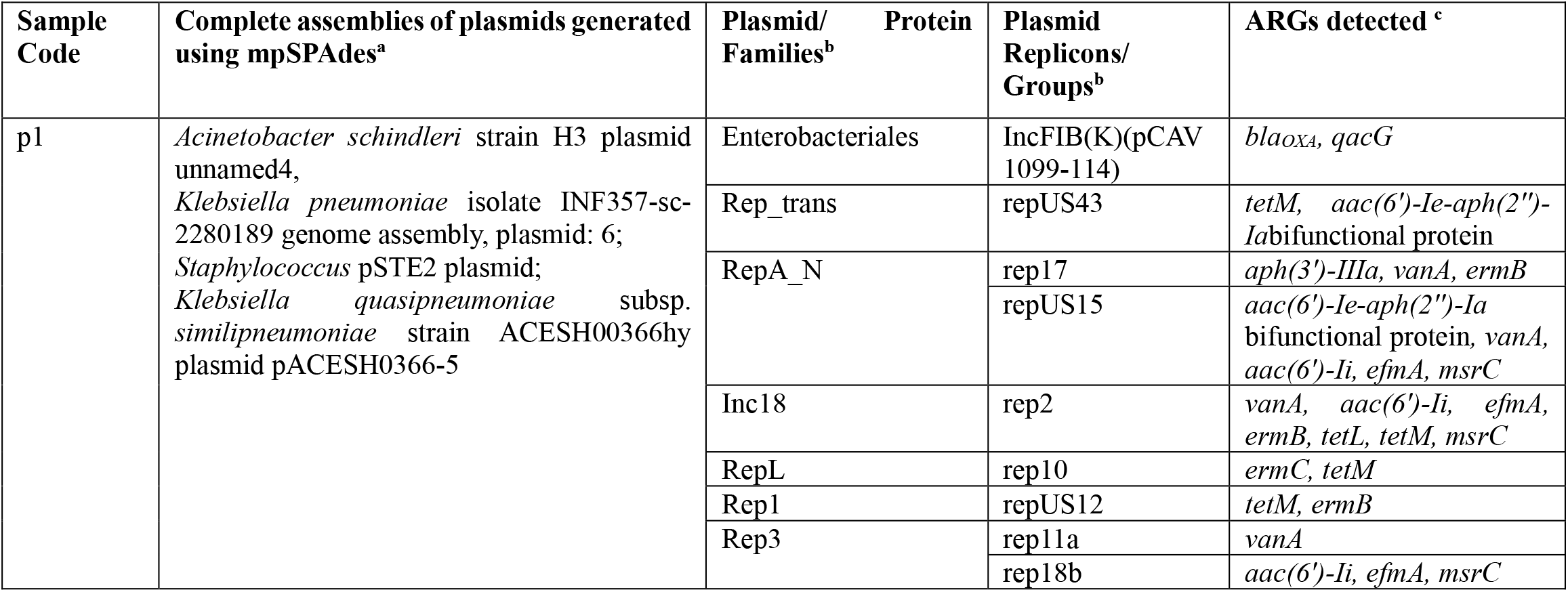
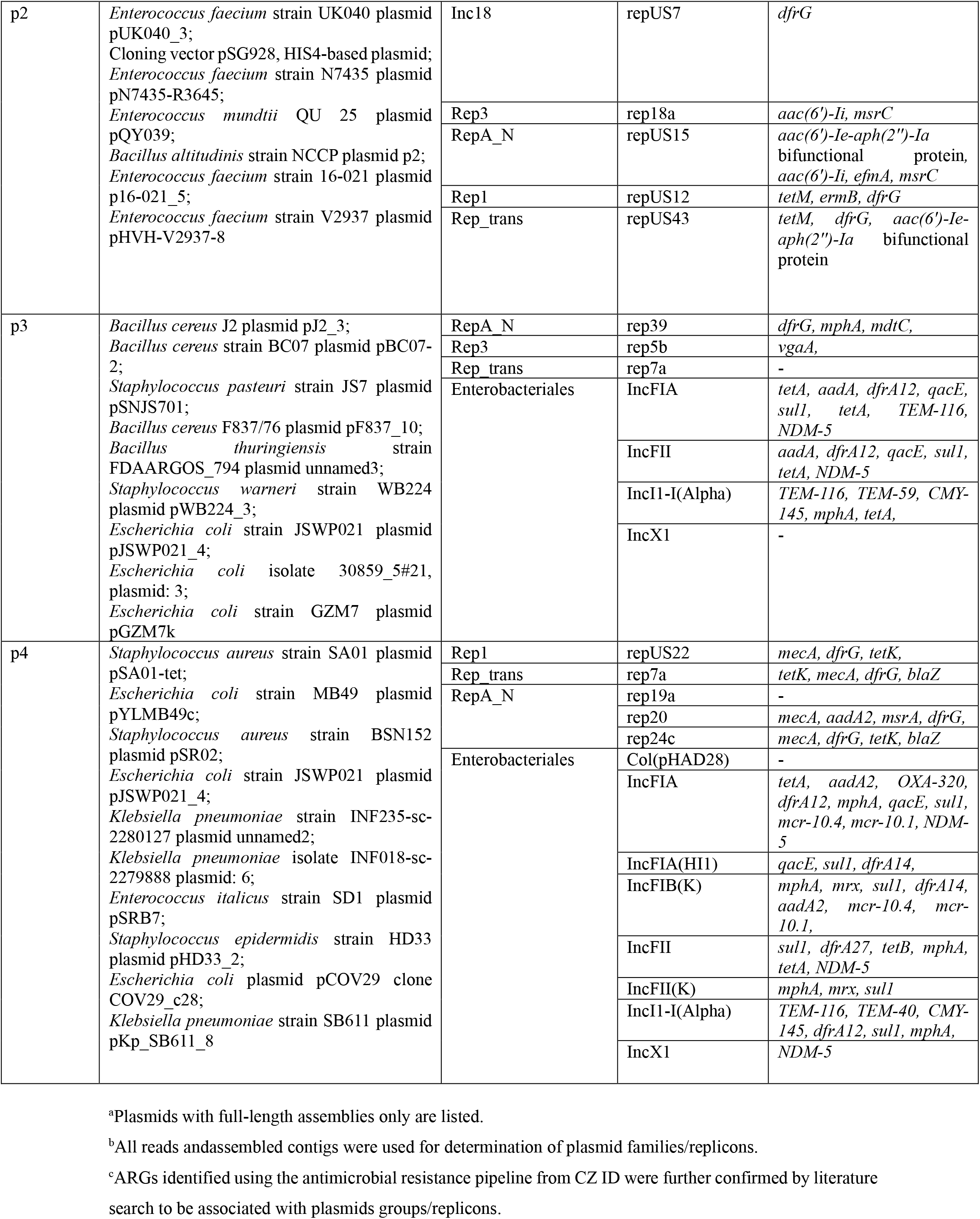
Identity of full-length Plasmids, Plasmid/ Protein Families, Plasmid Replicons and associated ARGs.

### 3.4 Resistome analysis – association of plasmid replicons and ARGs

Two independent resources, CARD and ResFinder, were used to detect and identify plasmid-associated ARGs. Using RGI-CARD, a total of 32, 26, 48 and 104 ARGs were detected in pDNA samples from p1, p2, p3 and p4 respectively (Table 4). A higher number of ARGs were identified using the RGI-CARD database as compared to ResFinder tool, thus highlighting the importance of using more than one database to identify the whole spectrum of potential ARGs in a sample. The ARGs belonged to diverse antibiotic/ drug classes like aminoglycosides, fluoroquinolones, tetracyclines, macrolides, nitroimidazole, carbapenems, monobactams, peptide antibiotics, glycopeptides, diamino pyrimidines, methicillin-resistant penams and sulfonamides, as well as to disinfecting agents and antiseptics. A summary of the ARGs detected per drug class is presented in Fig 3.

**Table 4.** Representative results of ARGs detected in the plasmid samples using RGI-CARD.

| Sample Code→ | p1 | p2 | p3 | p4 |
| --- | --- | --- | --- | --- |
| **Antibiotic mechanism**<br>↓ |  |  |  |  |
| **antibiotic efflux** | *kpnG, emrB, qacG, efmA, tetL, tetU, mtrD* | *mdtC, efmA, tetL, tetU, mdsC,* | *imrB, mdtC, mdtP, mdtO, mexQ, qacE, tetA* | *acrB, acrD, acrE, acrF, emrA, emrB, kpnG, kpnH, mdfA, mdtA, mdtC, msbA, cpxA, qacE, qacG, tetA, tetB, tetK, mexT* |
| **antibiotic inactivation** | *aac(6')-Ie-aph(2'')-Ia bifunctional protein, aac(6')-Ii, aph(3')-IIIa, OXA-360, OXA-277, bahA,* | *aac(6')-Ie-aph(2'')-Ia bifunctional protein, aac(6')-Ii, TEM-171, satA* | *aadA8b, CMY-145, mphA, mphB, mrx, NDM-5, TEM-116, TEM-59, satA, blaZ,* | *aadA8b, aadA2, ant(3'')-IIa, CMY-145, mphA, NDM-5, TEM-116, TEM-40, OXA-320, mrx, blaZ* |
| **antibiotic target alteration** | *ermB, ermC, vanA, vanH, vanR, vanX, vanZ,* | *ermB, ermT, rpsL, mupB, vanH, vanZ, vanY,* | *mupA, vanZ, vanT, vanW, vanY, ermZ* | *mcr-10.1, mcr-10.4, mupA, vanR, vanW, vanY* |
| **antibiotic target protection** | *msrC, tetM* | *msrC, tetM,* | *otrA, vgaA,* | *msrA,* |
| **antibiotic target replacement** | - | *dfrG* | *dfrA12, dfrG, sulI* | *dfrA12, dfrG, mecA, mecR1, sulI* |
| **reduced permeability to antibiotic** | *ompK37* | - | - | *ompA* |

**Fig. 3.**
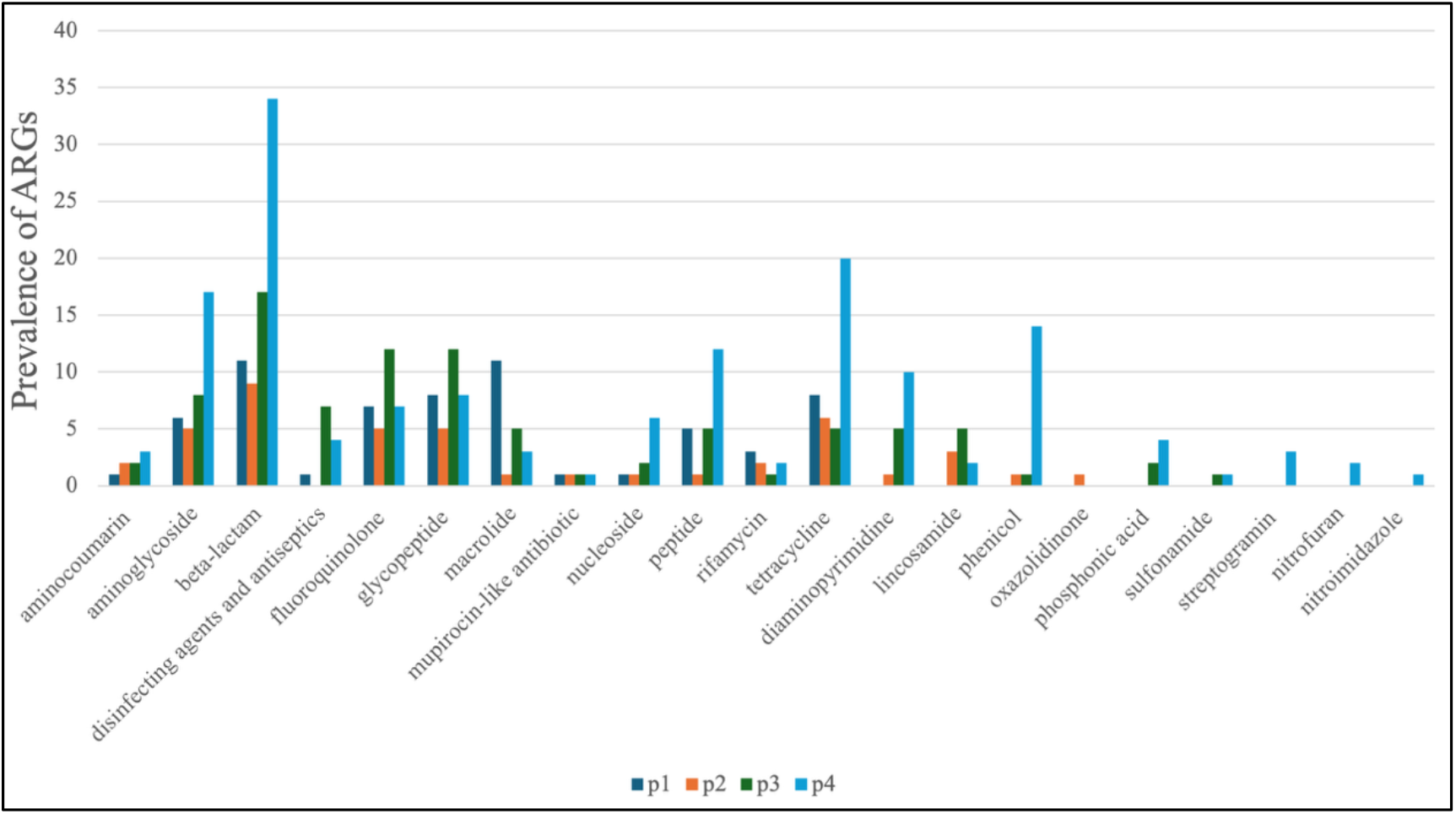
The graph depicts the relative abundance of different antibiotic classes in each of the four plasmid samples

## 4. Discussion

### 4.1 Prevalence of antibiotic-resistant bacteria in coastal water and their resistivity profile

Overall, the ARB load was found to be high in all water samples. Relatively higher bacterial CFU was observed in post-monsoon samples as compared to their pre-monsoon samples at the same location. This may stem from terrestrial wash offs where heavy monsoon rains flush land-based bacteria along with soil and debris into coastal waters. Agricultural fertilizers and organic matter also wash into the sea, providing a rich food source that helps heterotrophic bacteria multiply rapidly [6, 16]. Heavy rains frequently overwhelm urban drainage and sewage networks, leading to untreated waste discharges into water bodies, trigger phytoplankton growth thereby increasing overall bacterial count [17, 18].

### 4.2 Plasmid profile of MAR isolates

Though most isolates showed phenotypic resistance on AST, many did not give good plasmid yield upon extraction. Poor plasmid extraction from environmental bacterial isolates often stems from low copy number of wild type plasmids, change in the stress conditions of native bacterial environment vs lab conditions, shearing of the mega plasmids during extraction or presence of aggressive intracellular DNases. Other possibility could be that ARG(s) is/ are chromosomally located. However, since most isolates were multiple antibiotic resistant, the possibility of all ARGs being present on the bacterial chromosome is quite unlikely.

Subsequently, all four plasmid samples (p1, p2, p3 & p4) were processed individually for high throughput sequencing and analysis.

### 4.3 Plasmid profile of pre-monsoon samples

PlasmidFinder showed the presence of nine different types of plasmid replicons, namely IncFIB(K)(pCAV1099-114), repUS43, rep17, repUS15, rep2, rep10, repUS12, rep11a and rep18b in sample p1 (pre-monsoon Versova Beach, Table 3), and five different replicons like repUS7, rep18a, repUS15, repUS12 and repUS43 in sample p2 (pre-monsoon Juhu Beach). IncFIB(K)(pCAV1099-114) replicon is frequently found in *Klebsiella pneumonia* and *K. quasipneumoniae* [19, 20]. Other *Klebsiella*-associated plasmids detected in p1 were *Klebsiella pneumoniae* isolate INF357-sc-2280189 plasmid: 6 and *Klebsiella quasipneumoniae* subsp. *similipneumoniae* strain ACESH00366hy plasmid pACESH0366-5. Further, p1 sample detected plasmid replicons repUS43 & rep10, and plasmid pSTE2, all of which are known to be associated with *Staphylococcus* species [21]. Surprisingly, none of the isolates present in p1 sample were either *Klebsiella* sp. or *Staphylococcus* sp. This suggests the possibility of inter-species transfer of plasmids to isolates in p1sample. Taken with earlier studies [7, 22, 23], which show similar findings, it highlights inter-species transfer of plasmids by horizontal gene transfer as a major factor contributing to the spread of AMR in aquatic ecosystems.

The Inc18 incompatibility plasmids with replicons rep2 and repUS7 were detected in p1 and p2 samples respectively. Inc18 plasmids are most commonly found in vancomycin-resistant *Enterococcus* (VRE) isolates [24]. One of the isolates in each p1 and p2 samples belonged to VRE, indicating that the plasmids in these isolates might be carrying replicons rep2 and repUS7. VRE bacteria belong to the ESKAPEE group that comprises highly drug-resistant microbes [25]. IncFIB(K)(pCAV1099-114), detected in p1, is often found in *Klebsiella* sp. and carries *bla*_*OXA*_ (carbapenem resistance), one of the most prevalent and difficult-to-treat resistance [19, 26]. Transfer of such highly resistant traits to humans through recreational activities in coastal water can severely compromise clinical outcomes.

### 4.4 Plasmid profile of post-monsoon samples

Samples p3 and p4 (post-monsoon) detected nine and thirteen plasmids, respectively. Replicon analysis showed seven different types for p3 and of thirteen different types for p4. Various incompatibility (Inc) replicon groups were present in these samples (Table 3). Four incompatibility group of replicons, IncFIA, IncFII, IncI1-I(alpha), IncX1, present in p3, were also detected in p4. These plasmids are mostly associated with the *Enterobacteriaceae* family, mainly *E. coli*. Plasmid IncI family known to harbor genes for antibiotic resistance, have earlier been reported in *E. coli* isolated from poultry sources [27, 28], thus indicating inter-ecosystem exchange of resistant bacteria, most likely from animal husbandry waste flow into water bodies.

IncFIA plasmid replicons carry *bla*_*NDM*_ resistance gene that can easily transfer to other bacteria through different conjugative plasmids [29]. The different alleles of IncF detected in p3 and p4, namely, FII, FIA, FIB, are replicons which carry various resistance genes like ESBL and carbapenemases, genes encoding aminoglycoside-modifying enzymes, and plasmid-mediated quinolone resistance (PMQR) genes [30, 31]. Direct human contact with bacteria harbouring these replicons in recreational waters or through food chain via consumption of sea food can have widespread adverse implications on human health.

The rep7a and rep24c plasmid replicons detected in sample p4 are known to be associated with *mecA, blaZ, dfrG*, and *tetK*, conferring resistance to methicillin, carbapenems, trimethoprim and tetracyclines respectively [21]. Detection of these replicons and ARGs associated with them suggests possibility of their similar arrangement in plasmids of multidrug-resistant (MDR) isolates identified in the present study, which needs to be verified further.

The detection of diverse plasmid types and replicons, points to the heavy load and complexity of resistance factors and possibility of dynamic plasmid-based inter-species, inter-ecosystem ARG exchange, leading to rapid evolution and dissemination of AMR in water bodies.

### 4.5 Resistome analysis – association of plasmid replicons and ARG

The most frequent ARGs detected were the trimethoprim and aminoglycoside resistance genes *dfrG* and *vanZ*, that work by the mechanisms of target replacement and target alteration respectively. *dfrG* gene, detected in all samples except p1, is associated with various plasmid replicons like repUS7, repUS12, rep39, repUS22, rep7a, rep20, rep24c, etc. [21].

All samples were positive for various efflux pump based ARGs, known to give cross-resistance in bacteria, even to antibiotics belonging to different classes [32]. Their presence is concerning considering their potential for development of MDR strains in coastal water. Further analysis will reveal whether the efflux pumps are present individually or together on plasmids.

Beta-lactamase, which inactivates monobactams, was detected in p2 (*bla*_*TEM-171*_) gene, p3 and p4 (*bla*_*TEM-116*_). Variants of the *bla*_*OXA*_ carbapenem resistance gene (*OXA-360, OXA-277 and OXA-320*) were detected in samples p1and p4. These extended spectrum beta lactamases (ESBLs) are found in irrigation-purpose wastewaters and waste water treatment plants (WWTPs) [33, 34]. Their presence in the aquatic system highlights the probability of coastal waters receiving untreated/ partially treated waste water discharge from multiple sources.

Disinfecting agents and antiseptics resistance gene, the *qacG* gene, was detected in sample p1 and p4. *qacG* encodes for a multidrug resistance efflux pump that provides resistance to benzalkonium chloride, ethidium bromide and quaternary ammonium compounds (QACs). QACs are often associated with healthcare setups and have been found in clinical isolates [35]. Such biocide agents, despite being non-antibiotic in nature, can indirectly select for antibiotic resistance through co-selection, consequently increasing resistance to antibiotic drugs and other antimicrobial compounds, even in the absence of antibiotic selective pressure [1, 36]. Biocide resistance genes are more likely to be co-selected via co-resistance with ARGs. In-vitro experiments have demonstrated ability of biocides to co-select for antibiotic resistance in an aquatic bacterial community [37]. QAC resistant genes can be introduced into water bodies via multiple sources like municipal systems from household detergents, shampoos, surface cleaners, industrial products or untreated or partially treated WWTPs discharge [38, 39]. They have been found on plasmids containing ARGs for aminoglycoside resistance, beta-lactamase and trimethoprim, and on integrons of several Gram-negative and Gram-positive bacteria [40]. Thus, the presence of these genes on MGEs provides an increased likelihood of the resistance passing between environmental and clinically relevant bacterial strains [1]. Presence of the *qac* genes with such dynamic features not only points to their constant discharge into water bodies from various anthropogenic sources, but to a larger problem of AMR overload even in absence of antibiotic pressure.

The methicillin resistance *mecA* gene and *mecR1*gene, part of the BlaR1-MecR1regulatory complex leading to resistance to β-lactam antibiotics, were detected in sample p4 (post-monsoon Juhu Beach). Methicillin resistant *Staphylococcus aureus* (MRSA) is a leading cause of hospital and community-associated infections. It has inherent cross-resistance to several β-lactam antibiotics, one of the most widely used and effective classes of antimicrobials. Moreover, MRSA clinical strains being multi-drug resistant, therapeutic options for the treatment of *Staphylococcal* infections are often significantly hampered [41]. The concurrence of *qacG* gene, *mecA* gene and *mecR1 gene* and high prevalence of *qacG* gene in methicillin-resistant *Staphylococcal* species [35], strongly suggests their presence and co-selection in any or all three different *Staphylococcal* species identified in sample p4. Their probable release from hospital and pharmaceutical effluents to coastal water bodies can pose therapeutic challenge to public health.

The ARGs *tetM* and *ermC* were detected in p1 sample, and might be associated with repUS43 and rep10 plasmid replicons, as reported in earlier studies [21]. Besides *tetM*, repUS43 replicon is also known to carry the *aac(6’)-Ie-aph(2’’)-Ia* bifunctional protein gene which confers resistance to aminoglycoside antibiotics [42], again suggesting the presence of multiple ARGs on the same plasmid.

Genes involved in the regulation and expression of vancomycin resistance (*vanR, vanH, vanX, vanZ*, etc), frequently present in vancomycin-resistant *Enterococci* (VRE) species, were detected in all four samples. Gene *vanA* was identified only in p1 sample. Apart from *vanA* gene, ARGs *aac(6’)-Ie-aph(2’’)-Ia* and *aac(6’)-Ii*, both giving resistance to aminoglycosides, and *efmA, msrC* conferring resistance to macrolides were detected on same plasmid replicon repUS15. Acquisition of plasmids carrying multiple ARGs, belonging to different class of antibiotics, can give rise to MDR bacteria, untreatable by combination drug therapy, potentially compromising available treatment options [43].

A *Mycobacterium tuberculosis rpsL* gene (conferring resistance to Streptomycin) was detected in sample p2. However, none of the isolates from which sample p2 plasmid pool was prepared belonged to *M. tuberculosis*. Rather, plasmid DNA from *Micrococcus luteus* was present in sample p2. CARD contig_species hit also matched with *M. luteus*, suggesting that *M. luteus* most likely carried the *rpsL* gene, highlighting inter-species transfer of ARGs. Further characterization of *M. luteus* can confirm the presence of this gene on its pDNA.

Two variants of the *mcr* (mobile colistin resistance) gene, *mcr-10*.*4* and *mcr-10*.*1*, were detected in sample p4. The *mcr-10* gene was first identified in an IncFIA plasmid replicon and later in IncFIB [44], both replicons identified in p4 sample. Colistin is a last resort antibiotic used against carbapenem-resistant bacterial strains. Acquisition and dissemination of *mcr* genes to other non-colistin resistant strains in coastal environmental is therefore of great concern from clinical perspective.

The *bla*_*NDM-5*_ gene, conferring resistance to beta-lactam antibiotics (including last resort carbapenems), was found in 50% of the pDNA samples. The gene is easily transmissible and is associated with numerous incompatibility-type plasmid groups including the IncF, IncFIA, IncFII, IncFIB, IncX, [45], most of which were detected in sample p3 and p4. Recent report suggests that carbapenem-resistant *Salmonella enterica* serovar *typhi* (CR *Salmonella enterica* var *typhi*) strains carrying the *bla*_*NDM-5*_ gene have rapidly spread worldwide, especially across South Asia, including India and Pakistan, severely increasing treatment failures incidences and causing significant mortality. The strains exhibit extensive resistance to carbapenems, cephalosporins and fluoroquinolones, compromising the effectiveness of conventional therapeutic regimens [46]. Further in-depth studies to investigate the potential role of plasmids in *bla*_NDM-5_ dissemination in coastal waters is necessary.

A resistance gene, *mupB* was detected in p2 which confers strong resistance against mupirocin, a topical antibiotic used for the treatment of skin infections and eradication of MRSA strains [47]. Prevalence of *mupB* gene in clinical MRSA strains is known [48]. Interestingly, none of the isolates in sample p2 belonged to *S. aureus*, again indicating inter-species acquisition of this ARG. There are also no reports of its detection in coastal environments. Therefore, presence of *mupB* in coastal water highlights its probable source from hospital waste discharge and necessitates surveillance studies to monitor development of such rare clinical resistance in the aquatic environment. To the best of our knowledge, this is probably the first instance wherein plasmid mediated *mupB* gene has been detected in Arabian Sea coastal waters. The identification of such rare genes highlights the need for complementing robust surveillance technologies of water bodies based on high-throughput sequencing analysis with the culture-based methods.

The diversity of plasmid replicons and ARGs identified in the present study points to the complex nature of plasmid-associated ARGs. Their clinical relevance necessitates further investigation on impact of coastal water AMR on overall human health. The possibility of MDR strains in coastal waters can also lead to development of superbugs, which can disseminate globally. For example, ships lying idle for months due to various reasons including geo-political uncertainties, can develop extensive biofouling, which can carry invasive species to other oceanic areas when ships resume movement [49]. Although the present study cannot directly assign the identified ARGs to specific bacterial isolates or plasmids, since the extracted plasmid DNA was sequenced as pooled samples, it provides sufficient proof of various anthropogenic sources contributing to AMR in coastal water, like waste discharge from sewage, agriculture run-offs, domestic, industrial and hospital sources.

## 5. Conclusion

Mumbai, as the business capital of India, is a universal blueprint for land-sea pollution dynamics existing in most cities of the world with high international footprint. Moreover, as a major tourist hub, the local AMR conditions in Mumbai’s coastal waters can have global impact via consumption of contaminated seafood and recreational water activities. The resistance genes generated in Mumbai’s waters can also leave the area via international shipping (ballast water and biofouling), globalized seafood supply chains, or even migratory birds, thus having far-reaching global effects. Such local study data can be extrapolated to address broader AMR issues that apply worldwide. Hence, coordinated mitigation strategies are necessary at international level to address AMR effectively from global One Health Perspective.

## 7. Statements and Declarations

### Author Contributions

All authors contributed to the present study. Sampling, material preparation, experimentation and data consolidation was done by Devika Ghadigaonkar. Bioinformatics analysis was executed by Devashish Rath and Devika Ghadigaonkar. The conceptualization, designing and supervision of the study research work was carried out by Archana Rath. All authors have contributed towards writing and editing of the manuscript.

### Ethics approval and consent to participate

Not applicable as this research work did not involve human participants or animal subjects.

