## Supplementary data for "Using Arabian Sea Coastal Water as a Model to Study Plasmid-mediated AMR Dissemination driven by Anthropogenic Impacts"

**Title**

**Table S1.**

**Identity of bacterial isolates used in the study**

| **Isolate Code** | **Isolate identity** | **Location and Season of Sampling** | **Sample Code of pooled Plasmid DNA£** |
|---|---|---|---|
| 1 & 2 | *Enterococcus faecium* | Versova Beach, Pre-monsoon (latitude 19.130383° N and longitude 72.812931° E) | p1 |
| 3 | *Enterococcus viikkiensis* | | |
| 4 & 5 | *Micrococcus sp.* | | |
| 16 & 17 | *Bacillus sp.* | | |
| 6 | *Micrococcus luteus* | Juhu Beach, Pre-monsoon (latitude 19.10939° N and longitude 72.82407 ° E) | p2 |
| 7 & 8 | *E faecium* | | |
| 9 | *Enterococcus viikkiensis* | | |
| 10 | *Jonesia spp.* | | |
| 11 | *Micrococcus luteus* | | |
| 25 | *Enterobacter sp.* | | |
| 30 | *Bacillus sp.* | | |
| 13 | *Staphylococcus kloosii* | Versova Beach, Post-monsoon (latitude 19.130383° N and longitude 72.812931° E) | p3 |
| 14 | *Staphylococcus epidermidis* | | |
| 15 | *Bacillus sp.* | | |
| 18 | *Bacillus thuringiensis* | | |
| 19 & 20 | *Escherichia coli* | | |
| 21 | *Bacillus cereus* | | |
| 22 | *Micrococcus aloeverae* | Juhu Beach, Post-monsoon (latitude 19.10939° N and longitude 72.82407 ° E) | p4 |
| 23 | *Staphylococcus haemolyticus* | | |
| 24 | *Staphylococcus arlettae* | | |
| 26 | *Micrococcus sp.* | | |

| | |
|---|---|
| 27 | *Escherichia coli* |
| 28 | *Staphylococcus sp.* |
| 29 | *Bacillus sp.* |
| 31 | *Acinetobacter baumannii* |
| 32 | *Brachybacterium muris* |

**Molecular identification of the bacterial isolates was done using 16S rRNA gene PCR amplification following by Sanger sequencing and BLASTn analysis**

£- each sample type (p1-p4) contained pooled plasmids from the isolates belonging to that sample

**Table S2.**

**Antibiotic Sensitivity Test (AST) profile of selected MAR isolates to twelve clinically relevant antibiotics**

| Sample code | Isolate Code | Antibiotic → Isolate ↓ | AK | AZM | AT | CX | CIP | CTX | E | IMP | LE | TGC | TR | VA | MAR | MARi# |
|---|---|---|---|---|---|---|---|---|---|---|---|---|---|---|---|---|
| p1 | 1 | *Enterococcus faecium* | 23R | 13R | 6R | 19R | 30S | 15R | 15R | 30I | 23S | 25S | 6R | 14S | 7 | 0.58 |
| | 2 | *E. faecium* | 25R | 6R | 6R | 23R | 21S | 17R | 6R | 40I | 16S | 23S | 6R | 15S | 7 | 0.58 |
| | 3 | *E. viikkiensis* | 17R | 6R | 18R | 6R | 10R | 6R | 6R | 19R | 6R | 19R | 6R | 6R | 12 | 1.0 |
| | 4 | *Micrococcus sp.* | 22S | 6R | 6R | 20R | 17R | 17R | 6R | 34S | 14R | 22S | 6R | 15R | 9 | 0.75 |
| | 5 | *Micrococcus sp.* | 23S | 8R | 6R | 18R | 19R | 17R | 6R | 17R | 15R | 22S | 22S | 15R | 9 | 0.75 |
| | 16 | *Bacillus sp. (out of total 05 tested antibiotics)* | - | - | - | - | 23R | - | 22R | 32S | 22R | - | - | 12S | 3 | 0.6 |
| | 17 | *Bacillus sp. (out of total 05 tested antibiotics)* | - | - | - | - | 25I | - | 20R | 30S | 21R | - | - | 12S | 2 | 0.4 |
| p2 | 6 | *M.luteus* | 18S | 20R | 6R | 15R | 21R | 15R | 24S | 30S | 22R | 21S | 6R | 15R | 8 | 0.67 |
| | 7 | *E. faecium* | 20R | 17R | 6R | 17R | 24S | 16R | 26R | 34I | 21S | 21S | 6R | 12S | 7 | 0.58 |
| | 8 | *E. faecium* | 20R | 13R | 6R | 14R | 17S | 18R | 21R | 25I | 16S | 21S | 6R | 16S | 7 | 0.58 |
| | 9 | *E. viikkiensis* | 22R | 19R | 6R | 25R | 27S | 20R | 25R | 28I | 25S | 21S | 18R | 13S | 7 | 0.58 |
| | 10 | *Jonesia spp.* | 17S | 6R | 13R | 6R | 10R | 6R | 6R | 17R | 8R | 21S | 6R | 6R | 10 | 0.83 |
| | 11 | *Micrococcus luteus* | 16S | 6R | 14R | 6R | 11R | 6R | 6R | 16R | 7R | 21S | 6R | 6R | 10 | 0.83 |
| | 25 | *Enterobacter sp.* | 18S | 12S | 25I | 20S | 27S | 25S | 12R | 22S | 23S | 20S | 23S | 6R | 2 | 0.17 |
| | 30 | *Bacillus sp. (out of total 05 tested antibiotics)* | - | - | - | - | 28I | - | 30S | 32S | 27I | - | - | 6R | 1 | 0.2 |

| | | | | | | | | | | | | | | | |
|---|---|---|---|---|---|---|---|---|---|---|---|---|---|---|---|
| p3 | 13 | *Staphylococcus kloosii* | 22S | 20R | 6R | 24S | 26I | 25S | 24S | 36S | 24I | 22S | 21S | 13R | 3 | 0.25 |
| | 14 | *Staphylococcus epidermidis* | 22S | 21S | 6R | 26S | 31I | 19R | 25S | 37S | 27I | 23S | 20S | 12R | 3 | 0.25 |
| | 15 | *Bacillus sp. (out of total 05 tested antibiotics)* | - | - | - | - | 21R | - | 19R | 33S | 22R | - | - | 14S | 3 | 0.6 |
| | 18 | *Bacillus thuringiensis(out of total 05 tested antibiotics)* | - | - | - | - | 19R | - | 18R | 29R | 20R | - | - | 12S | 4 | 0.8 |
| | 19 | *Escherichia coli* | 21S | 8R | 9R | 6R | 8R | 6R | 6R | 14R | 8R | 20S | 7R | 6R | 10 | 0.83 |
| | 20 | *Escherichia coli* | 21S | 10R | 15R | 6R | 7R | 6R | 6R | 13R | 6R | 18S | 6R | 6R | 10 | 0.83 |
| | 21 | *Bacillus cereus (out of total 05 tested antibiotics)* | - | - | - | - | 24I | - | 25S | 30S | 22R | - | - | 13S | 1 | 0.2 |
| p4 | 22 | *Micrococcus aloeverae* | 27S | 22S | 16R | 27S | 25I | 25S | 29S | 29S | 23I | 26S | 27S | 6R | 2 | 0.17 |
| | 23 | *Staphylococcus haemolyticus* | 27S | 17R | 14R | 28S | 18R | 27S | 22S | 17R | 18R | 26S | 20S | 18R | 6 | 0.5 |
| | 24 | *Staphylococcus arlettae* | 29S | 16R | 6R | 23S | 24I | 18R | 22S | 32S | 24S | 25S | 6R | 15R | 5 | 0.42 |
| | 26 | *Micrococcus sp.* | 16S | 11R | 24R | 18R | 24I | 26S | 8R | 18R | 24I | 18R | 20S | 6R | 7 | 0.58 |
| | 27 | *Escherichia coli* | 20S | 10R | 15R | 6R | 8R | 6R | 6R | 12R | 8R | 20S | 6R | 6R | 10 | 0.83 |
| | 28 | *Staphylococcus sp.* | 30S | 10R | 6R | 26S | 34I | 20R | 20R | 34S | 33S | 25S | 6R | 15R | 6 | 0.5 |
| | 29 | *Bacillus sp. (out of total 05 tested antibiotics)* | - | - | - | - | 23R | - | 24S | 30S | 21R | - | - | 12S | 2 | 0.4 |
| | 31 | *Acinetobacter baumannii* | 18R | 20R | 12R | 9R | 20R | 11R | 20R | 22I | 20R | 19R | 8R | 6R | 11 | 0.92 |
| | 32 | *Brachybacterium muris* | 27S | 22S | 16R | 6R | 27I | 6R | 22S | 39I | 24I | 23S | 6R | 23R | 5 | 0.42 |

Abbreviations: AK = amikacin; AZM = azithromycin; AT = aztreonam; CX = cefoxitin; CIP = ciprofloxacin; CTX = cefotaxime; E = erythromycin; IPM = imipenem; LE = levofloxacin; TGC = tigecycline; TR = trimethoprim; VA = vancomycin; R = resistant; S = sensitive; I = intermediate; MAR = multiple antibiotic resistance; MARi = multiple antibiotic resistance index

Colour coding: red = resistant; green = sensitive; black = intermediate

MAR: Multiple Antibiotic Resistant

[#]MARi: MAR out of the total antibiotics used for that isolate

Numbers indicate zone of inhibition (ZOI) in millimeter (mm)

Results have been compared using the EUCAST break points table, 2022

**Fig S1.**

**Representative plasmid DNA profile of select MAR isolates.**

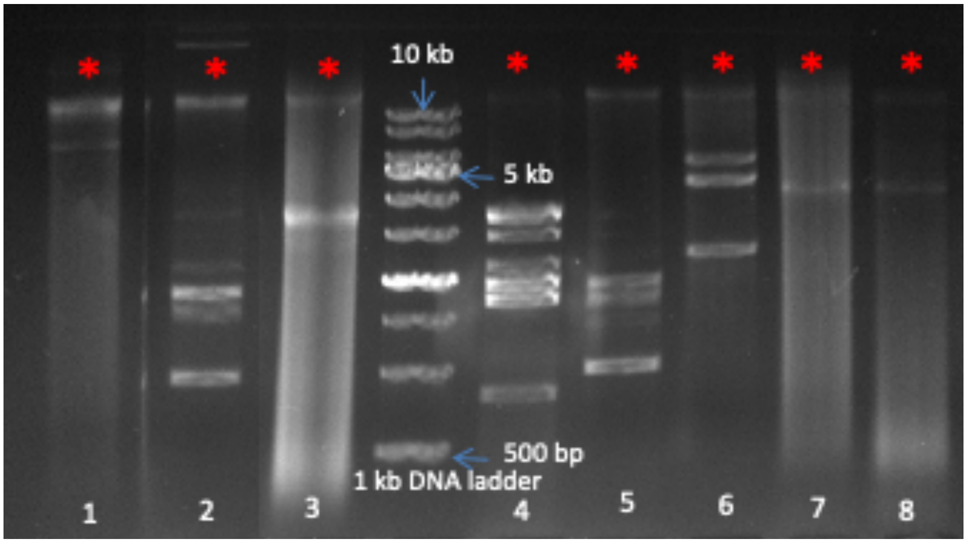

**All isolates revealed presence of multiple plasmids including high molecular weight plasmids (> 10 kb; marked *)**

Lanes 1-8: 1: *Micrococcus aloeverae*; 2: *Staphylococcus arlettae*; 3: *Acinetobacter baumannii*; 4: *Enterobacter* spp.; 5: *Escherichia coli*,; 6: *Staphylococcus kloosii*; 7: *Bacillus* spp.; 8: *Bacillus thuringiensis*

**Table S3.**
**Details of the plasmid sequences deposited on NCBI's GenBank and their corresponding accession numbers**

| Sample Code | Sequence Name | Sequence Identity from BLASTn | Accession number |
|---|---|---|---|
| p1 | NODE_1_length_6453_type_circular | *Acinetobacter schindleri* strain H3 plasmid unnamed4 | PZ221391 |
| | NODE_2_length_3824_type_circular | *Klebsiella pneumoniae* isolate INF357-sc-2280189 genome assembly, plasmid: 6 | PZ221392 |
| | NODE_3_length_2600_type_circular | *Staphylococcus* pSTE2 plasmid | PZ221393 |
| | NODE_4_length_2425_type_circular | *Klebsiella quasipneumoniae subsp. similipneumoniae* strain ACESH00366hy plasmid pACESH0366-5, complete sequence | PZ221394 |
| p2 | NODE_1_length_11765_type_circular | *Enterococcus faecium* strain UK040 plasmid pUK040_3, complete sequence | PZ221395 |
| | NODE_1_length_3127_type_circular | Cloning vector pSG928, HIS4-based plasmid, complete sequence | PZ221396 |
| | NODE_2_length_58766_type_circular | *Enterococcus faecium* strain N7435 plasmid pN7435-R3645, complete sequence | PZ221397 |
| | NODE_3_length_27251_type_circular | *Enterococcus mundtii* QU 25 plasmid pQY039 DNA, complete genome | PZ221398 |
| | NODE_4_length_7927_type_circular | *Bacillus altitudinis* strain NCCP plasmid p2, complete sequence | PZ221399 |
| | NODE_5_length_3991_type_circular | *Enterococcus faecium* strain 16-021 plasmid p16-021_5, complete sequence | PZ221400 |
| | NODE_6_length_2183_type_circular | *Enterococcus faecium* strain V2937 plasmid pHVH-V2937-8, complete sequence | PZ221401 |
| p3 | NODE_1_length_19291_type_circular | *Bacillus cereus* J2 plasmid pJ2_3 DNA, complete sequence | PZ221402 |
| | NODE_1_length_58736_type_circular | *Bacillus cereus* strain BC07 plasmid pBC07-2, complete sequence | PZ221403 |
| | NODE_2_length_10136_type_circular | *Staphylococcus pasteuri* strain JS7 plasmid pSNJS701, complete sequence | PZ221404 |
| | NODE_3_length_8462_type_circular | *Bacillus cereus* F837/76 plasmid pF837_10, complete sequence | PZ221405 |
| | NODE_4_length_7340_type_circular | *Bacillus thuringiensis* strain FDAARGOS_794 plasmid unnamed3, complete sequence | PZ221406 |
| | NODE_5_length_4422_type_circular | *Staphylococcus warneri* strain WB224 plasmid pWB224_3, complete sequence | PZ221407 |
| | NODE_6_length_4208_type_circular | *Escherichia coli* strain JSWP021 plasmid pJSWP021_4, complete sequence | PZ221408 |
| | NODE_7_length_3422_type_circular | *Escherichia coli* isolate 30859_5#21 genome assembly, plasmid: 3 | PZ221409 |
| | NODE_8_length_1678_type_circular | *Escherichia coli* strain GZM7 plasmid pGZM7k, complete sequence | PZ221410 |
| p4 | NODE_1_length_13945_type_circular | *Staphylococcus aureus* strain SA01 plasmid pSA01-tet, complete sequence | PZ221411 |

| | | | |
|---|---|---|---|
| | NODE_1_length_41856_type_circular | *Escherichia coli* strain MB49 plasmid pYLMB49c, complete sequence | PZ221412 |
| | NODE_3_length_22381_type_circular | *Staphylococcus aureus* strain BSN152 plasmid pSR02 | PZ221413 |
| | NODE_4_length_4208_type_circular | *Escherichia coli* strain JSWP021 plasmid pJSWP021_4, complete sequence | PZ221414 |
| | NODE_5_length_4018_type_circular | *Klebsiella pneumoniae* strain INF235-sc-2280127 plasmid unnamed2, complete sequence | PZ221415 |
| | NODE_6_length_3063_type_circular | *Klebsiella pneumoniae* isolate INF018-sc-2279888 genome assembly, plasmid: 6 | PZ221416 |
| | NODE_7_length_3059_type_circular | *Enterococcus italicus* strain SD1 plasmid pSRB7, complete sequence | PZ221417 |
| | NODE_8_length_2552_type_circular | *Staphylococcus epidermidis* strain HD33 plasmid pHD33_2 | PZ221418 |
| | NODE_9_length_1678_type_circular | *Escherichia coli* plasmid pCOV29 clone COV29_c28 | PZ221419 |
| | NODE_10_length_1367_type_circular | *Klebsiella pneumoniae* strain SB611 plasmid pKp_SB611_8, complete sequence | PZ221420 |

*The raw data is available on NCBI's Sequence Read Archive (SRA) with BioProject accession number PRJNA1422493*
